# The mushroom body output encodes behavioral decision during sensory-motor transformation

**DOI:** 10.1101/2022.09.14.507924

**Authors:** Cansu Arican, Felix Johannes Schmitt, Wolfgang Rössler, Martin Fritz Strube-Bloss, Martin Paul Nawrot

## Abstract

Animal behavioral decisions are dynamically formed by evaluating momentary sensory evidence on the background of individual experience and the acute motivational state. In insects, the mushroom body (MB) has been implicated in forming associative memories and in assessing the appetitive or aversive valence of sensory stimuli to bias approach versus avoidance behavior. To study the MB involvement in innate feeding behavior we performed extracellular single-unit recordings from MB output neurons (MBONs) while simultaneously monitoring a defined feeding behavior in response to timed odor stimulation in naïve cockroaches. All animals expressed the feeding behavior almost exclusively in response to food odors. Likewise, MBON responses were invariably and strongly tuned to the same odors. Importantly, MBON responses were restricted to behaviorally responded trials, which allowed the accurate prediction of the occurrence versus non-occurrence of the feeding behavior in individual trials from the neuronal population activity. During responded trials the neuronal activity generally preceded the onset of the feeding behavior, indicating a causal relation. Our results contest the predominant view that MBONs encode stimulus valence. Rather, we conclude that the MB output dynamically encodes the behavioral decision to inform downstream motor networks.

## Introduction

Animal behavioral decisions are based on the processing of momentary environmental conditions on the background of innate and experience dependent behavioral biases. In insects, and specifically in nocturnal species, olfactory cues play a major role in a variety of behavioral decisions involved e.g. in mating, oviposition, or navigation. During foraging, locating food sources and evaluating their quality is fundamental for survival and requires the accurate recognition of appropriate food odors to inform behavioral decisions.

Feeding behavior in insects is experimentally accessible through the registration of well-defined behaviors, e.g. in the proboscis extension response (PER) in bees (Bitterman et al., 1983) and flies (Yetman and Pollack, 1987; Shiraiwa and Carlson, 2007) and the maxilla-labia response (MLR) in cockroaches (Arican et al., 2020). The individual decision of an animal about executing a feeding behavior can be modulated by its internal and behavioral states (see Discussion). These influences may enter at different stages of sensory-motor processing including the mushroom body (MB) (Devineni and Scaplen, 2022).

While olfactory processing and learning in insects is being studied in great detail, we still lack understanding of how and at which stage of the recurrent sensory-motor pathway behavioral decisions are formed. Here we take advantage of the experimental accessibility in the cockroach that allows us to simultaneously monitor feeding behavior and record from central brain neurons in the individual animal with high temporal resolution. The MB is an evolutionary old and homologous central brain structure in insects (Strausfeld et al., 2009). The MB output integrates sensory input of different modalities (Li and Strausfeld, 1999; Yagi et al., 2016; Strube-Bloss and Rössler, 2018) through sensory projections with the internal state, the behavioral state and external sensory context (Cohn et al., 2015; Tsao et al., 2018; Siju et al., 2020; Aimon et al., 2022) through a large number of recurrent, mostly neuromodulatory input (see Discussion). While the MB function has predominantly been assigned to the formation and recall of short and long term associative memories (Heisenberg, 2003; Menzel, 2012; Hige et al., 2015; Owald et al., 2015), recent studies in untrained animals have demonstrated an important role of the MB in processing attractive and repulsive sensory stimuli in the context of innate behaviors (Bräcker et al., 2013; Lewis et al., 2015; Tsao et al., 2018; Siju et al., 2020) as well as its involvement in state-dependent sensory-motor transformation (Okada et al., 1999; Aimon et al., 2022).

Previous studies, mostly conducted in honey bees (Strube-Bloss et al., 2011, 2016) and fruit flies (Aso et al., 2014b; Owald and Waddell, 2015; Hancock et al., 2022), have provided accumulated evidence that distinct populations of MB output neurons (MBONs) establish a code for the valence of a sensory stimulus with respect to its behavioral relevance and specifically so as a consequence of associative learning (see Discussion). Here we demonstrate that a subpopulation of MBONs in the cockroach faithfully predicts the occurrence or non-occurrence of a defined feeding behavior in the cockroach on a single trial basis. Our results therefore contest the prevailing view that the output of the MB merely encodes the valence of sensory stimuli and we conclude instead that, at its output, the MB represents an integrated signal of internal state, momentary environmental conditions and experience-dependent memory to encode a behavioral decision.

## Results

### Feeding behavior is expressed in response to food odors

We start out with analyzing the odor specific feeding behavior in individual animals. We presented a set of 10 different odors and one additional control stimulus (clean air, Fig. 1) to each animal. Each stimulus is repeated 10 times (trials). The continuous recording of the animals’ mouthparts allowed for the detection of a behavioral feeding response on a trial-by-trial basis. For each of the 10 trials per odor we thus obtained a binary data (no MLR vs. MLR) on the animal’s feeding response.

**Figure 1.**
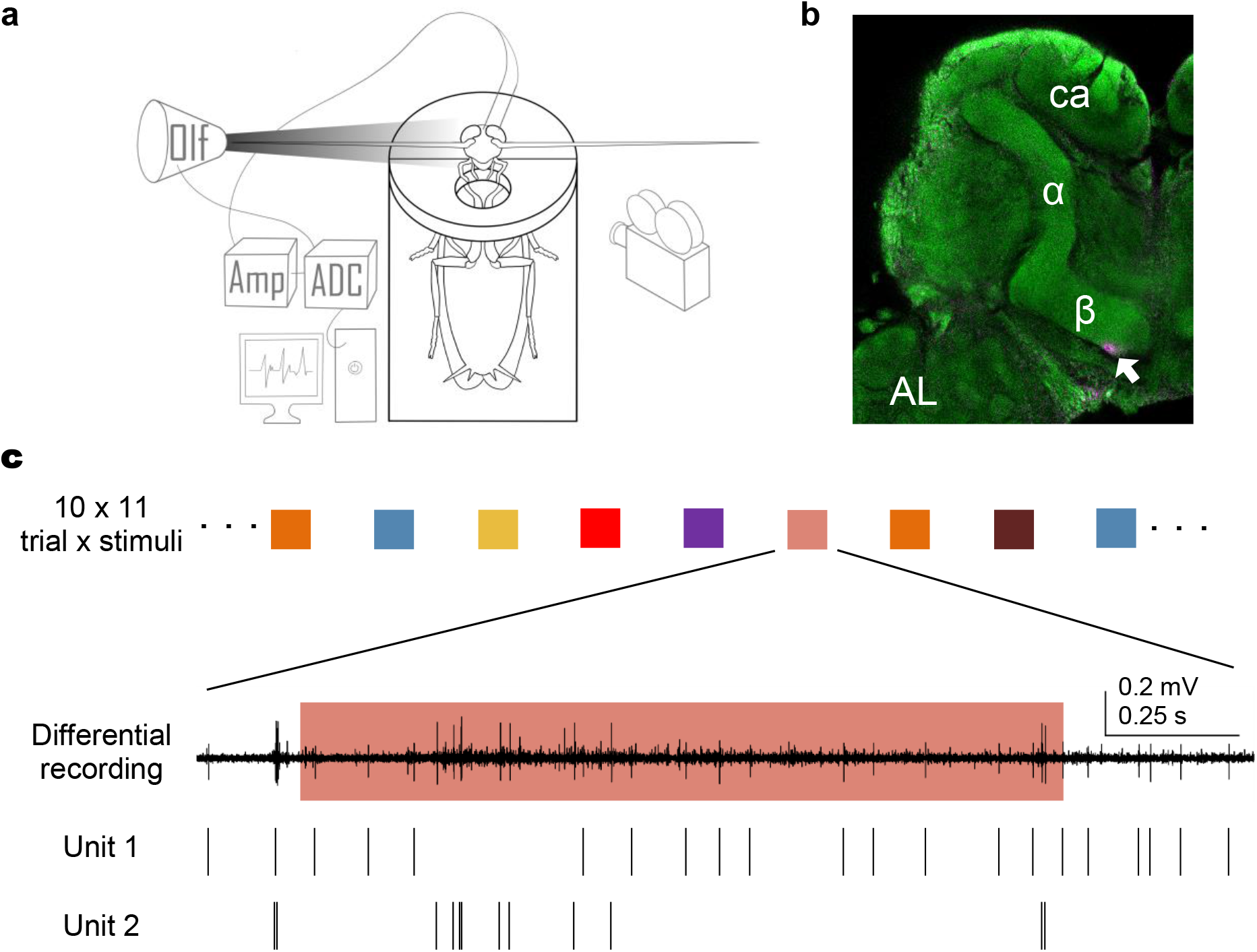
Extracellular single-unit recordings from MB output with simultaneous recording of feeding behavior. **a** Schematic illustration of a harnessed cockroach with inserted electrodes connected to an amplifier (Amp) and analog to digital converter (ADC). A computer-controlled odor supply system (Olf) provides the odor stimulus to the antenna from its tip. A camera records the mouthparts of the cockroach allowing for the detection of the maxilla-labia response (MLR). **b** Background staining with Lucifer Yellow of the right hemisphere of a cockroach brain. The MB α and β lobes are clearly visible. The position of the electrode tip, shown in magenta, (white arrow) is located at the border of the MB β lobe where the primary neurites of the MBONs leave the MB. AL: antennal lobe, ca: mushroom body calyx. **c** Stimulation pattern with odor and control stimulations in pseudo-randomized order. Extracellular differential recording during a single stimulation of 2 s duration (light red shading) with the odor cinnamaldehyde and the corresponding spiking activity of two single units.

We observed a clear overall behavioral response pattern across the set of tested odors in the group of animals (Fig. 2a). Only a small subset of four odors triggered repeated feeding behavior. In contrast, and consistently across all animals, almost no responses were evoked by the presentation of the remaining six odors. The few sparsely distributed responses to this subset of odors matches the low spontaneous response probability to the control stimulus (overall 2.2%). Importantly, the three responded odors isoamyl acetate (Iso), cinnamaldehyde (Cin) and benzaldehyde (Ben) are known food odors in the cockroach (Arican et al., 2020; Khoobdel et al., 2021).

**Figure 2.**
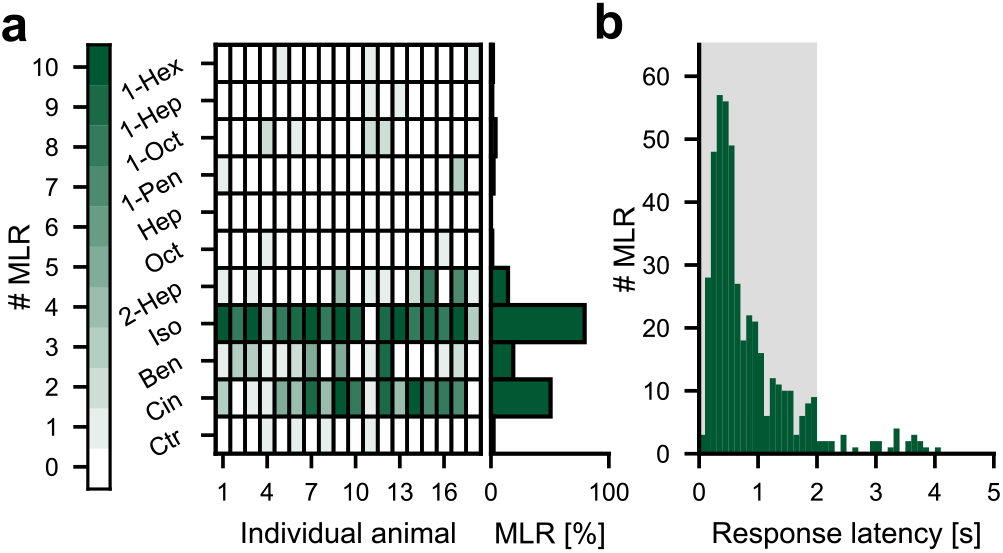
Behavioral feeding response exclusively occurs to food odors. **a** Number of observed feeding responses (MLR) over 10 trials per stimulus for the control and all odors in individual animals (matrix) and average across all animals (horizontal bar histogram) during the 2 s odor presentations. Stimuli: 1-hexanol (1-Hex), 1- heptanol (1-Hep), 1-octanol (1-Oct), 1-pentanol (1-Pen), heptanal (Hep), octanal (Oct), 2-heptanone (2-Hep), isoamyl acetate (Iso), benzaldehyde (Ben), trans-cinnamaldehyde (Cin), control (Ctr). **b** Distribution of single trial behavioral response latency during and after odor presentation (binwidth = 100 ms), gray shaded area depicts stimulus presentation. The mean (median) behavioral response time was 776 ms (632 ms).

### Neuronal population response is dominated by food odors

To gain insight into the role of the MB output in sensory-motor transformation we recorded extracellular single unit activity from the output region of the MB β lobe throughout the experiment (Fig. 1). In a first analysis we quantified the single neuron and population activity in response to odor stimulation. Across the MBON population we observed clear and consistent responses only to the three food odors that also evoked consistent behavioral responses (Fig. 3d). In Fig. 3a we show, as an example, the single-trial spike trains of a single MBON and the corresponding trial-averaged firing rates in Fig. 3b. This neuron shows a strong response to Iso, a weak response to Cin and no response to Ben. Analysis of the trial-averaged neuronal population rate in Fig. 3c shows different population response profiles for the three odors. The fraction of responding neurons was largest for Iso and smallest for Ben, reflecting the behavioral odor response pattern in Fig. 2a. The control stimulus did not evoke a response in any neuron.

**Figure 3.**
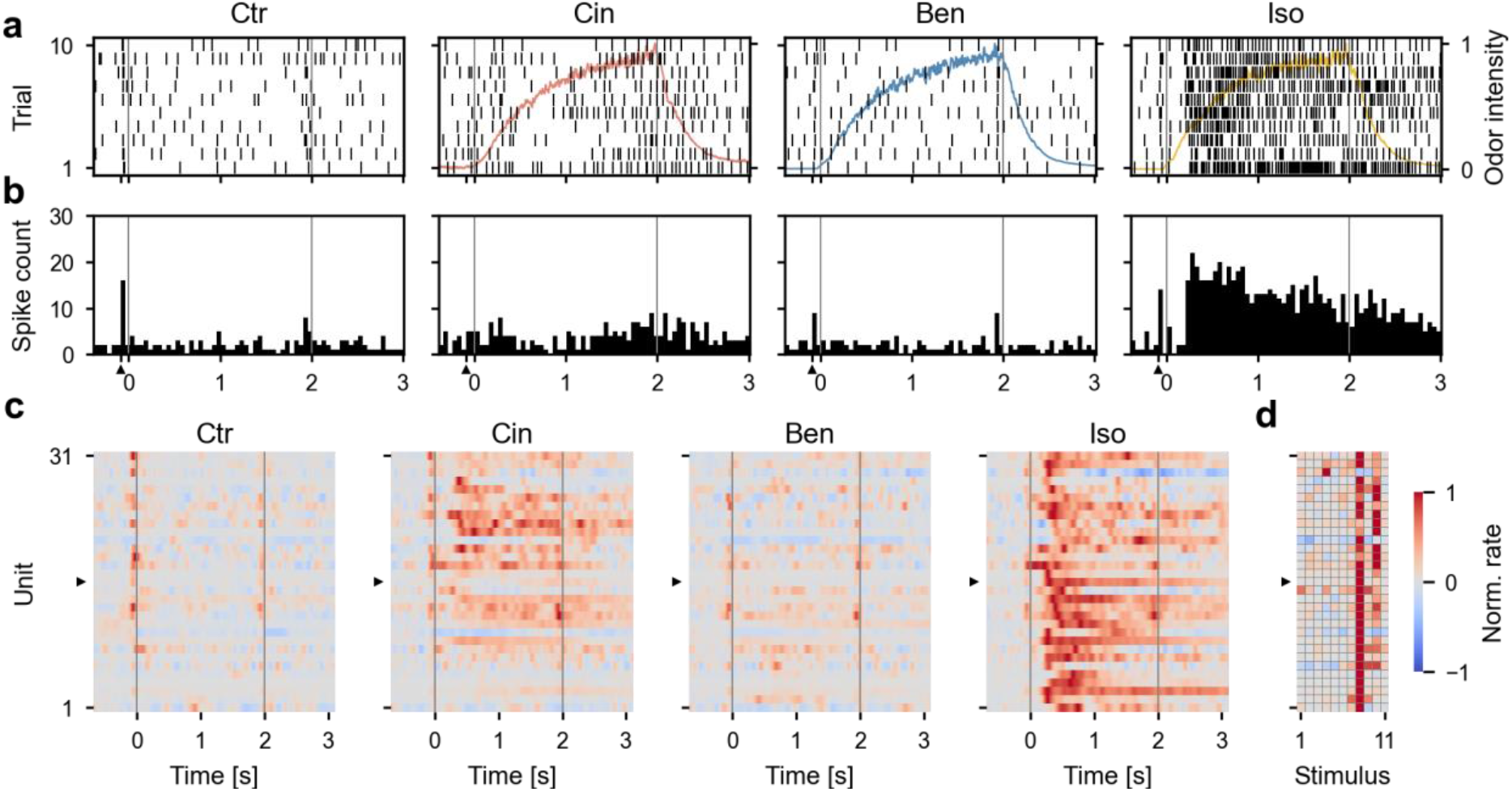
Neuronal population response during odor stimulation. **a** Exemplary spike raster plots of unit 16 (animal 10) during 10 repeated presentations of four different stimuli (Ctr = control, Cin = cinnamaldehyde, Ben = benzaldehyde, Iso = isoamyl acetate). The normalized trial-averaged odor concentration as measured with the PID is shown in the background (light color). **b** Peristimulus time histograms across all 10 stimulus trials (binwidth = 50 ms). Upward pointing triangle (▲) depicts the time point of valve switch at t = −90 ms before the odor arrives at the antenna tip (t = 0 ms). **c** Normalized trial-averaged firing rates across the population of 31 neurons. Color code indicates changes in the normalized firing rates relative to the baseline firing rate. **d** Matrix depicts normalized time-averaged firing rates across all units and all stimuli during stimulus onset (order of odors left to right as in Fig. 2a top to bottom). Odor stimulus was presented in the interval 0 – 2 s. Rightward pointing triangle (▸) in c & d indicates unit 16 shown in a & b.

### MBON responses reflect behavioral feeding responses to food odors

We now consider the relation of single neuron responses and behavioral responses to the three food odors (Cin, Ben, Iso). To this end, and for each animal, we sorted all trials in two groups with respect to their behavioral response (MLR vs. no MLR). As a first result we find that the response spike count was consistently higher in behaviorally responded trials than in behaviorally unresponded trials for all recorded neurons (Fig. 4a).

**Figure 4.**
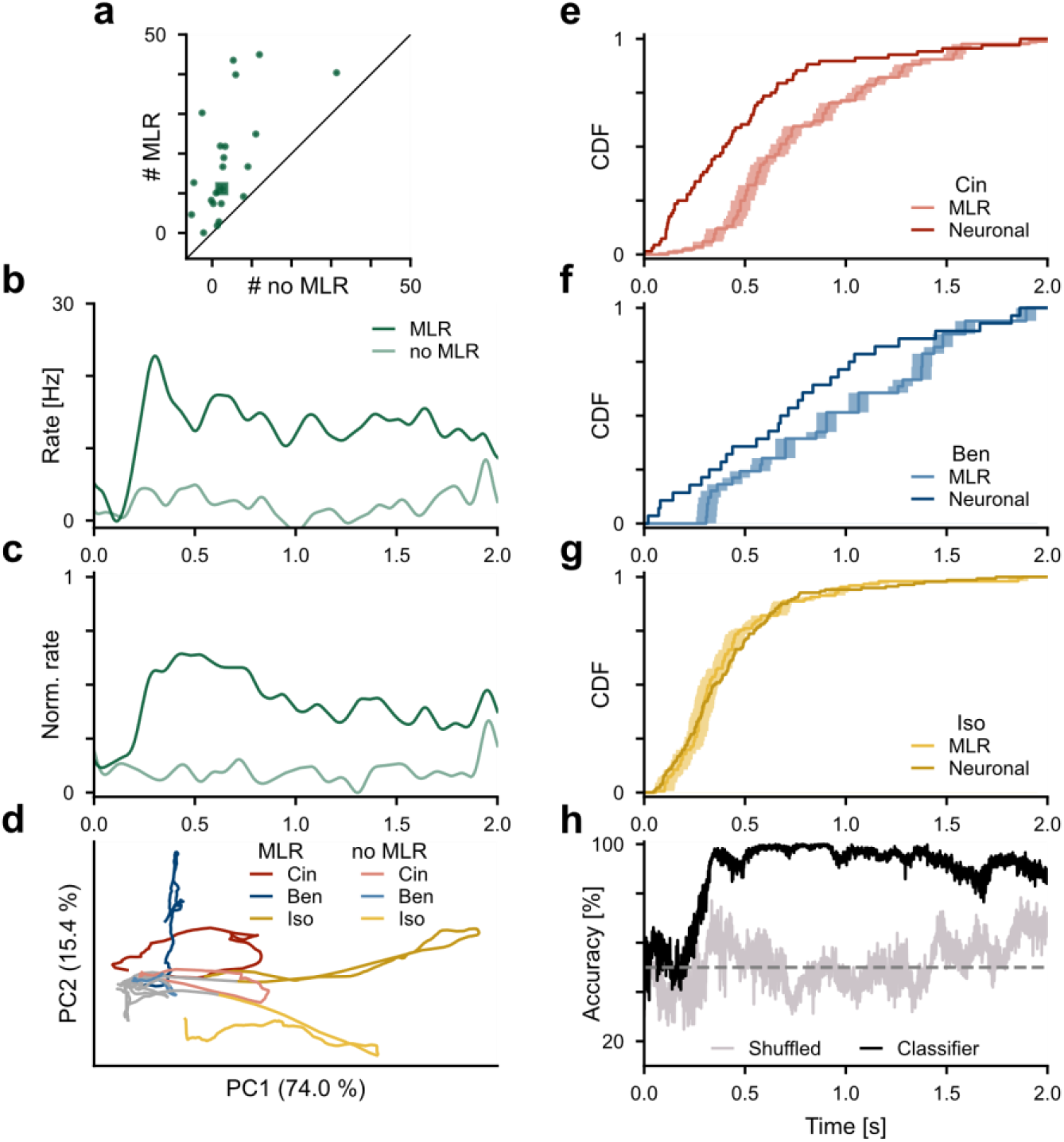
Mushroom body output reliably predicts behavioral response to food odors. **a** The median spike count during presentation of the food odors (Cin, Ben, Iso) computed across trials and per neuron is larger in behaviorally responded trials (MLR, ordinate) than in behaviorally unresponded trials (no MLR, abscissa) in all MBONs. **b** The time-resolved trial-averaged firing rate of a single MBON (unit 15, animal 10) shows a strong response to stimulation with food odors in behaviorally responded (MLR) but not in unresponded (no MLR) trials. **c** Across neuron average (N = 23) of normalized firing rate responses to food odors sorted by the animals’ behavior indicate a clear neuronal response only to behaviorally responded trials but none to behaviorally unresponded trials. **d** First against the second principal component (PC1, PC2) of MBON population response separated by stimulus (Cin, Ben, Iso) and by behavioral response (MLR vs. no MLR) before (gray) and during (colored) stimulus presentation indicates separability of odors and behavioral state. **e-g** Across the single-trial MLR (weak line; shading refers to the 100 ms duration of the individual video frame) and single-trial neuronal response onset time (strong line) during presentation of Cin (e), Ben (f) and Iso (g). Only trials, which exhibit a behavioral response are considered (MLR). **h** Accuracy of single-trial prediction of behavioral response from neuronal population activity (black) sharply increases ∼250 ms after stimulus onset and reaches almost perfect prediction. Averaged single trial prediction accuracy of behavioral response (MLR) based on a logistic regression classifier. The test data set (black) is compared to a control data set with randomly shuffled MLR labels (gray).

Next, we analyzed the time-resolved response rate for the two behavioral conditions. The exemplary firing rate profile of a single MBON in Fig. 4b demonstrates a clear and strong trial-averaged response during trials that lead to a behavioral response, while no response occurred during behaviorally unresponded trials. Averaging the normalized firing rates across all MBONs confirms this picture where behaviorally responded trials show a prominent increase in firing rate in response to the olfactory stimuli whereas behaviorally unresponded trials did not evoke a discernible response (Fig. 4c), indicating a differential involvement of MBONs in the two behaviorally distinct categories.

To study in more detail the neuronal representation of odor and behavioral category we performed a principal component analysis (PCA) on the MBON population. Considering the 1^st^ and 2^nd^ principal components in Fig. 4d, we find that for each of the three odors, the respective trajectory during behaviorally responded trials (MLR) separates from the trajectory during behaviorally unresponded trials (no MLR), confirming our result on the averaged neuronal responses in Fig. 4c. In line with previous results in the honey bee (Strube-Bloss et al., 2011) we also observe a distinct representation of all three food odors in the principal component space.

### Neuronal responses precede behavioral responses

What is the relative timing of behavioral and neuronal response? To answer this question, we estimated behavioral and neuronal response latencies at the single-trial level. In each behaviorally responded trial we determined the first video frame after odor onset (t = 0) that indicated a MLR. The overall distribution of behavioral response latencies shown in Fig. 2b is skewed towards the time of odor onset. In Fig. 4e-g we compare, per odor, the cumulative distributions of neuronal and behavioral single trial responses. While for Cin (Fig. 4e) and Ben (Fig. 4f) the neuronal responses are detected earlier than the behavioral responses, the situation is more difficult to assess for Iso (Fig. 4g) where neuronal and behavioral responses were overall fastest.

Note, that we applied a conservative strategy for estimating the single trial neuronal response onset, which minimizes the possibility of a premature response detection. At the same time and due to the stochastic nature of spike responses this approach tends to overestimate single trial latencies, specifically in weakly responded trials. To alleviate this estimation bias we used an additional approach where for each neuron we estimated its response latency from the trial averaged firing rate across all behavioral response trials for a given odor. This resulted in average single unit response latencies of 353 ms (range: 40-1,658 ms), 577 ms (67-1,821 ms) and 208 ms (27-1,575 ms) for the three odors Cin, Ben and Iso, respectively, indicating that individual MBONs express different response latencies.

### Neuronal population activity allows for the single-trial prediction of the behavioral response

Can we decode the neuronal response to predict feeding behavior in the single trial? To answer this question, we trained and tested a machine learning classifier in a time-resolved manner. To this end we first labeled the single trial neuronal responses according to their behavioral outcome (class label MLR vs. no MLR). We then separated the complete data set in a training and a test set. The former was used to train the classification algorithm in order to predict the class label based on the single-trial neuronal population activity. Classification performance was then evaluated on the test data set and quantified as accuracy. In a cross-validation approach we repeated the procedure on different splits of training and test data sets. The result in Fig. 4h (black curve) shows the average accuracy as a function of trial time. Initially, after stimulus onset, accuracy is at chance level (50%, horizontal dashed line) before it rises sharply to plateau at a high level of >90% accuracy. As a control we repeated the complete classification approach on the same data set, albeit with randomly shuffled behavioral class labels. As a result, accuracy did not significantly deviate from chance level.

## Discussion

Over the past decade, a series of experimental studies on associative olfactory conditioning have concluded that the MB output encodes the valence of a sensory stimulus (see Introduction). The large majority of studies have been conducted in the fruit fly *Drosophila melanogaster*. With few exceptions (e.g. Sayin et al., 2019; Siju et al., 2020), these experiments evaluated learning induced plasticity at the level of MBONs and behavioral memory expression during a memory retention test in a behavioral group assay (Tully and Quinn, 1985) in separate groups of animals that underwent the same classical conditioning protocol (Aso et al., 2014b; Hancock et al., 2022). This approach did not allow to match neuronal and behavioral responses in the same individual and on a trial-to-trial basis.

By taking advantage of the experimental accessibility in the cockroach that allows us to simultaneously record neuronal spiking activity and feeding behavior with high temporal resolution in the individual animal, we were in the position to perform combined trial-based analyses of neuronal and behavioral responses. Our results demonstrate a tight link between the neuronal response at the MB output and the actual execution of a defined feeding behavior on a trial-to-trial basis where the occurrence or non-occurrence of behavior could be faithfully predicted with a single-trial classification approach. Our results contest the predominant view that the MB merely encodes sensory stimulus valence that provides a stimulus dependent behavioral preference. From our data we conclude that the MB output momentarily encodes a behavioral decision that is required for the execution of a behavior. Indeed, it has been shown that MBONs project to premotor areas (Li and Strausfeld, 1997, 1999; Okada et al., 2003; Aso et al., 2014a) and it has recently been shown that they can also make direct connections to descending neurons that innervate the ventral nerve cord (Hsu and Bhandawat, 2016; Emanuel et al., 2020; Li et al., 2020) in fruit flies and cockroaches.

Our conclusion is in line with earlier experimental observations in the cockroach (Mizunami et al., 1998; Okada et al., 1999) and recent experimental interpretations in the fruit fly (Hige et al., 2015; Lewis et al., 2015; Tsao et al., 2018; Siju et al., 2020; Aimon et al., 2022), which have suggested a tighter and acute involvement of the MB output in motor control. By investigating innate behavior in the fruit fly, Tsao et al. (2018) conclusively showed that MBON output is required for the expression of food-seeking behavior. This result implies that MBON activity should causally precede the behavioral execution. With respect to the precise relative timing of MBON activity and behavioral response, our analyses show that, on average, the onset of physiological spiking responses in the recorded MBON population indeed preceded the feeding behavior of the mouth parts. Stimulus response latencies differed across individual MBONs where several neurons showed fast stimulus-response times as short as 40 ms while others show considerably late response onsets after several hundred milliseconds and thus during the actual behavior. This feature of MBON specific response latencies matches earlier results in MBON recordings from naïve honey bees that reported a considerable fraction of fast responding MBONs that establish a rapid encoding of odor identity within ∼70-80 ms (Strube-Bloss et al., 2012). Interestingly, we find that both, neuronal and behavioral response latencies are odor specific. Isoamyl acetate, the major single molecule component of banana blend (Schubert et al., 2014) and highly attractive both for flies and cockroaches (Schubert et al., 2014; Arican et al., 2020; Khoobdel et al., 2021), provoked the fastest and strongest neuronal as well as the fastest behavioral responses. We may further suggest that the later MBON responses occurring during behavior reflect on the behavioral state of the animal and rely on feedback signals (Mizunami et al., 1998; Okada et al., 1999). Indeed, in the fruit fly it has been shown that ongoing walking behavior through feedback via dopaminergic, octopaminergic and serotonergic neuromodulatory neurons strongly can dynamically influences MB activity (Cohn et al., 2015; Siju et al., 2020; Aimon et al., 2022).

In summary we hypothesize that the MB lobes are positioned at the center of the sensory-motor loop where it continuously integrates sensory evidence and monitors the animal’s metabolic and current behavioral state to form behavioral decision that is encoded in the MB output.

## Materials and Methods

### Animals

For all experiments adult male *Periplaneta americana* were used. Laboratory colonies were kept at 26 °C with a reversed light-dark cycle (12 h: 12 h) and fed with oat flakes and water *ad libitum*. All experiments were conducted during the scotophase, the natural active phase of *P. americana*.

### Experimental setup

For data acquisition an extracellular recording setup was used (Fig. 1). The recording electrode (adapted from Okada et al., 1999; Strube-Bloss et al., 2011) consisted of three polyurethane coated copper wires (Ø 14 µm; Electrisola, Escholzmatt, Switzerland) glued together with hard sticky wax (Siladent, Goslar, Germany). A Teflon coated silver wire (Ø 125 µm, World Precision Instruments) was used as reference. Wires were fixed to a head stage that was connected to a preamplifier (PA 103, Electronics Workshop, University of Cologne, Germany). Main amplification using a 4-channel amplifier (MA 102 differential amplifier, Electronics Workshop, University of Cologne, Germany) was performed in differential mode from all three possible pair combinations of electrode wires. Amplified signals were bandpass filtered (300 Hz to 5 kHz), A/D converted with 16-bit amplitude resolution and a sampling rate of 25 kHz using a data acquisition unit (CED Micro 1401 mk II, Cambridge, UK) and stored on a PC.

An odor supply system (adapted from Strube-Bloss et al. (2011) was customized (by the workshop of the Department of Biology and an electrical engineer of the Institute of Zoology at the University of Cologne, Germany). The air stream (3.5 LPS charcoal filtered air) was split into two pathways. The first provided a permanent airstream (regulated with a restrictor, ∅ = 0.25 mm). The second passed through 12 magnetic valves (LFAA1200118H, Lee, Sulzbach, Germany). Each valve enables computer-controlled switching to pass the airstream either through an empty glass bottle (100 ml volume) or through a glass bottle (100 ml volume) filled with 5 ml of a pure odorant. The air outlet was placed at the tip of the right antenna aligned to its longitudinal axes such that the complete antenna was covered by the airstream (Fig. 1a).

A video camera (Logitech QuickCam Pro) was positioned in front of the animal’s head to capture movement of its mouthparts, enabling the detection of a behavioral response (MLR, Arican et al., 2020).

All parts of the setup were placed in a Faraday cage covered in opaque fabric for shielding. Experiments were conducted in red light (643 nm), invisible to the animals (Goldsmith and Ruck, 1958; Walther, 1958; Mote and Goldsmith, 1970).

### Animal preparation

Animals were anesthetized by cooling to 4 °C and harnessed in a custom-made holder (Fig. 1). The neck was fixed with hard sticky wax. Maxillary palps and the first basal segment of the antennae (scapus) were fixed with periphery wax (Sigma Dental Surgident, Systems, Handewitt, Germany) to avoid movement and contact with the electrodes. A small window was cut between the compound eyes and above the bases of the antennae. Trachea and head glands on top of the brain were removed and the animal was placed in the Faraday cage with a magnetic stand. Visible contours of the brain were used as landmarks to place the electrode above the β lobe in the right brain hemisphere, the silver wire was placed in the left compound eye. The recording electrode was slowly inserted axially with a micromanipulator (Luigs & Neumann SM-6, Ratingen, Germany) while monitoring the recording signal until the typical large amplitude extracellular action potentials of MBONs were detected. Then the head was covered with periphery wax to avoid dehydration of the brain and electrode movement.

### Stimulus protocol

Ten odors we used for stimulation: 1-hexanol (Merck KGaA, Darmstadt, Germany), 1-heptanol (Merck KGaA, Darmstadt, Germany), 1-octanol (Thermo Fisher Scientific, Waltham, MA, USA), 1-pentanol (Merck KGaA, Darmstadt, Germany), heptanal (Thermo Fisher Scientific, Waltham, MA, USA), octanal (Merck KGaA, Darmstadt, Germany), 2-heptanone (Thermo Fisher Scientific, Waltham, MA, USA), isoamyl acetate (Thermo Fisher Scientific, Waltham, MA, USA), benzaldehyde (Merck KGaA, Darmstadt, Germany) & trans-cinnamaldehyde (Merck KGaA, Darmstadt, Germany). This set of odors is composed of single molecular odors that are perceivable by cockroaches (Sakura et al., 2002; Arican et al., 2020; Paoli et al., 2020; unpublished data). The food odors isoamyl acetate (Iso), benzaldehyde (Ben) and trans-cinnamaldehyde (Cin) were included because they are known to be able to elicit feeding behavior in cockroaches (Arican et al., 2020) and are clearly attributable to the specific food sources banana, almond, and cinnamon, respectively. The stimulation protocol comprised 10 trials per odor and additional 10 trials for the control stimulus (clean air). The total of 110 stimulations were presented in a different pseudorandomized order to each animal. The odors were presented for 2 s with an inter-trial interval (ITI) of 30 s (Fig. 1c).

To control for stimulus timing, we performed separate calibration measurements where odor concentration was measured at the antennal tip and base using a high temporal resolution photoionization detector (200B miniPID, Aurora Scientific, Aurora, ON, Canada). Odors consistently arrived at the antennal tip position with a delay of 90 ms with respect to the time of valve switching. We defined this time point as stimulus onset for subsequent analyses. In addition to the odor response some neurons showed a very brief spiking response that occured with high temporal precision immediately after valve switching and before the odor arrived at the antennal tip (Fig. 3b, ▲). We argue that switching a valve between the clean and the odorous air-stream caused a very brief interruption of the air stream detectable as a brief mechanical stimulus. We therefore interpret this initial brief and accurately timed neuronal response as a mechanosensory response. In addition, visual stimulation with either ultraviolet or bluish green light generated responses in a large subset of neurons (not shown) reflecting the MB role in multisensory integration.

### Data processing and spike sorting

Preprocessing of extracellular recordings was performed with the software Spike 2 (v7.2, CED, Cambridge, UK). Semi-automated spike detection and a template-based spike sorting algorithm (Spike 2) was used. For spike detection we defined a threshold of minimum three times the signal’s standard deviation as computed outside the stimulation intervals. Templates of the spike waveform were generated by the software and subsequently manually revised. Simultaneous video recordings (10 fps with constant frame rate) were made with the Spike 2 Video Recorder (v1.05, CED, Cambridge, UK) and analyzed manually in Spike 2 for MLR detection. In total, recordings of 21 animals were analyzed. From 19 animals 31 single units could be extracted and the MLR could be analyzed in 18 animals. In 15 animals neuronal signals and behavior were analyzed simultaneously.

### Visualization of recording position

To visualize the recording tract, the tip of the electrode was dipped in Alexa Fluor 647 Hydrazide (Invitrogen, Thermo Fisher Scientific, Waltham, MA, USA) before insertion into the brain. After recording, the brain was dissected in fresh cockroach Ringer’s solution (185 mM NaCl, 4 mM KCL, 6 mM CaCl2, 2 mM MgCl2, 10 mM Hepes, 35 mM Glucose, pH 7.2) and the whole brain was moved to 4% Formaldehyde in 0.1 M PBS overnight on a shaker at 4 °C. On the next day, the tissue was washed with 0.1 M PBS and 0.2% Triton X-100 in 0.1 M PBS with each solution three times for 10 min. Afterwards, the brain was incubated in 0.5% Lucifer Yellow CH dilithium salt (Merck KGaA, Darmstadt, Germany) diluted in 0.1 M PBS overnight on a shaker at 4 °C and washed again with 0.1 M PBS (5x 10 min) the next day. Finally, the brain was dehydrated in ethanol (30%, 50%, 70%, 90%, 95%, 2x 100%; 10 min each step), cleared in methylsalycylate (VWR Chemicals, Radnor, PA, USA) and mounted in it. The recording tract was visualized using a confocal laser scanning microscope (Leica TCS SP8, Wetzlar, Germany) and the scans were processed with ImageJ (FIJI based on ImageJ 1.53c, Wayne Rasband, NIH, USA).

### Firing rate estimation

For analysis and visualization of neural and behavioral data custom written code in Python 3 was used. Firing rates were estimated by kernel convolution (Nawrot et al., 1999; Meier et al., 2008) with a time resolution of 1 ms. Trials were first aligned to stimulus onset (t = 0). Four different kernel functions were used: (1) a symmetric non-causal (i.e. centered) Gaussian kernel 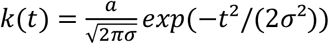 supported on [−3*σ*, 3*σ*], (2) an asymmetric and strictly causal exponential kernel *k*(*t*) = *a · exp*(−*t*/*τ*) supported on [0, 5*τ*], (3) an asymmetric non-causal alpha-shaped kernel (Krofczik et al., 2008) 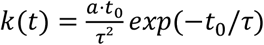 for *t*_0_ > 0 and 0 otherwise aligned to its center of gravity (*t*_0_ = *t* + 1.6783 *· τ*) and supported on [−5*τ*, 5*τ*], (4) a strictly causal alpha-shaped kernel aligned at *t*_0_ = *t* supported on [0, 6*τ*]. All kernels were normalized to unity such that 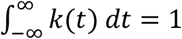.

For each neuron rates were estimated from single trials and from the pooled responses across trials.

To compute the baseline-corrected trial-averaged firing rate responses to odor stimulation per neurons we pooled spike trains across the respective set of trials and performed convolution with a Gaussian kernel (*σ* = 50 ms) and divided the result by the number of trials. We then subtracted the baseline firing rate as averaged across the baseline time window [−20 s, −0.5 s].

To obtain normalized firing rates, we averaged the firing rates per neuron and divided them by the maximum firing rate of the respective neuron over all odors.

### Principal component analysis

For PCA we selected units from animals that showed a behavioral response (MLR) to food odors (Cin, Ben, Iso) during at least one trial and at most during nine trials per odor such allowing for comparison between behaviorally responded and unresponded trials. Six animals (seven neurons) fulfilled this condition. Single trial rate estimates (non-causal alpha kernel, *τ* = 200 *ms*) were used. The mean firing rate in the baseline window [−20 *s*, −0.5 *s*] was subtracted in each single trial. Single trial rates were grouped by odor identity and behavioral outcome (MLR vs. no MLR) before trial-averaging. All vectors were used together for PCA.

### Response latencies

To determine single trial neuronal response latencies, we used the baseline uncorrected data. Latencies were estimated from rate estimates with the causal exponential kernel (*τ* = 250 *ms*). For this, baseline activity was estimated in the time window [−20 *s*, −0.5 *s*]. For each trial the 97-percentile of the baseline firing rate in the time window [−20 *s*, −0.5 *s*] was determined and used as threshold. If this value was smaller than 2.25 *· max*(*k*(*t*)) the threshold was set to 2.25 *· max*(*k*(*t*)). Response latency was determined as the time point of the first threshold crossing of the response rate from below within the time window [0 *s*, 2 *s*]. This conservative procedure minimizes the chance to detect any premature response caused by spontaneous activity. As a consequence, there is a tendency to overestimate single trial response latencies when a response is detected only after integrating several true response spikes.

Based on the estimated firing rates of the single-trial analysis population onsets were analyzed. The single trials were grouped by their odor and unit identity. Baseline and stimulus activity was averaged across all trials in these groups. For each group the 97-percentile of the baseline firing rate was determined as response threshold. Neuronal onsets were detected in the group firing rate stimulus activity of the group, if a value was greater than the threshold as the border. If a neuronal onset was detected, the time point of the first crossing from below was considered as the time point of the neuronal onset. The mean across all units per odor was estimated. Units without a detected onset were ignored.

### Prediction of behavioral outcome

For single trial prediction of a behavioral response from population activity during stimulation with food odors (Cin, Ben, Iso) we employed logistic regression classification with L2-norm (Python package Scikit-learn, Pedregosa et al., 2011) with regularization strength of 1.0. For training and testing the classifier we constructed neuronal pseudo populations (Rickert et al., 2009) as follows. We first selected those animals that expressed feeding behavior (MLR) in at least 10 and at most 20 out of total 30 stimulation trials resulting in a population of 17 units. We then estimated single trial firing rates using a causal alpha kernel (*τ* = 50 *ms*) and subtracted the mean firing rate in the baseline window [−20 *s*, −0.5 *s*]. All trials were grouped into two behavioral classes (MLR vs. no MLR). For each neuron and both classes, trials were then split into test (67%) and training set (33%). Training and testing were performed for each time point separately by randomly drawing (with replacement) 50 training and 20 test samples such that each sample consisted of a vector of 17 single trial firing rates. The mean and standard deviation of the training samples was estimated. Training and test data was centered by this mean and scaled by this standard deviation. Prediction accuracy was calculated as the percentage of correctly predicted single trial behavioral outcomes in the test set. The training and test procedure was repeated 24 times based on independent random splits into test and training sets. As a means of control, the complete process is repeated for a surrogate data set with randomly shuffled MLR labels, predicting a performance at chancel level (50%).

## Data Availability

The data sets generated for this study are available on request to the corresponding authors.

## Authors contributions

CA and MN designed the research. CA conducted the experiments. CA and FS analyzed the data. CA and MN wrote the manuscript. MS-B supported the adaptation of the recording setup. MS-B and WR revised the manuscript.

## Conflict of interest statement

The authors declare that the research was conducted in the absence of any commercial or financial relationships that could be construed as a potential conflict of interest.

## Acknowledgements

We thank Claudia Groh for sharing and teaching the staining protocol for the electrode tract. We thank Michael Dübbert of the electronics workshop of the Neurophysiology section of the Institute of Zoology for providing us with solutions for the electrophysiological setup and the computer-controlled odor stimulation. We thank the workshop of the Department of Biology headed by Leo Lesson for constructing the odor supply system. We thank Vivian Theling for support in establishing the staining protocol in our lab. This project was funded by DFG-FOR 2705 (grant no. 365082554). CA and FS received a PhD scholarship from the Research Training Group Neural Circuit Analysis on the Cellular and Subcellular Level funded through the German Research Foundation (DFG-GRK 1960, grant no. 233886668 to MN).

## Notes

### Competing Interest Statement

The authors have declared no competing interest.

### Summary of Updates

The name of one author and a funding number were corrected.

## References

Aimon, S., Cheng, K. Y., Gjorgjieva, J., and Grunwald Kadow, I. C. (2022). Walking elicits global brain activity in Drosophila. bioRxiv, 1–38. doi:10.1101/2022.01.17.476660.

Arican, C., Bulk, J., Deisig, N., and Nawrot, M. P. (2020). Cockroaches show individuality in learning and memory during classical and operant conditioning. Front. Physiol. 10, 1–14. doi:10.3389/fphys.2019.01539.

Aso, Y., Hattori, D., Yu, Y., Johnston, R. M., Iyer, N. A., Ngo, T.-T., et al. (2014a). The neuronal architecture of the mushroom body provides a logic for associative learning. Elife 3, 1–47. doi:10.7554/eLife.04577.

Aso, Y., Sitaraman, D., Ichinose, T., Kaun, K. R., Vogt, K., Belliart-Guérin, G., et al. (2014b). Mushroom body output neurons encode valence and guide memory-based action selection in Drosophila. Elife 3, 1–42. doi:10.7554/eLife.04580.

Bitterman, M. E., Menzel, R., Fietz, A., and Schäfer, S. (1983). Classical conditioning of proboscis extension in honeybees (Apis mellifera). J. Comp. Psychol. 97, 107–119. doi:10.1037/0735-7036.97.2.107.

Bräcker, L. B., Siju, K. P., Varela, N., Aso, Y., Zhang, M., Hein, I., et al. (2013). Essential role of the mushroom body in context-dependent CO2 avoidance in Drosophila. Curr. Biol. 23, 1228–1234. doi:10.1016/j.cub.2013.05.029.

Cohn, R., Morantte, I., and Ruta, V. (2015). Coordinated and compartmentalized neuromodulation shapes sensory processing in Drosophila. Cell 163, 1742–1755. doi:10.1016/j.cell.2015.11.019.

Devineni, A. V., and Scaplen, K. M. (2022). Neural circuits underlying behavioral flexibility: insights from Drosophila. Front. Behav. Neurosci. 15, 1–24. doi:10.3389/fnbeh.2021.821680.

Emanuel, S., Kaiser, M., Pflueger, H.-J., and Libersat, F. (2020). On the role of the head ganglia in posture and walking in insects. Front. Physiol. 11, 1–11. doi:10.3389/fphys.2020.00135.

Goldsmith, T. H., and Ruck, P. R. (1958). The spectral sensitivities of the dorsal ocelli of cockroaches and honeybees. J. Gen. Physiol. 41, 1171–1185. doi:10.1085/jgp.41.6.1171.

Hancock, C. E., Rostami, V., Rachad, E. Y., Deimel, S. H., Nawrot, M. P., and Fiala, A. (2022). Visualization of learning-induced synaptic plasticity in output neurons of the Drosophila mushroom body γ-lobe. Sci. Rep. 12, 10421. doi:10.1038/s41598-022-14413-5.

Heisenberg, M. (2003). Mushroom body memoir: from maps to models. Nat. Rev. Neurosci. 4, 266–275. doi:10.1038/nrn1074.

Hige, T., Aso, Y., Rubin, G. M., and Turner, G. C. (2015). Plasticity-driven individualization of olfactory coding in mushroom body output neurons. Nature 526, 258–262. doi:10.1038/nature15396.

Hsu, C. T., and Bhandawat, V. (2016). Organization of descending neurons in Drosophila melanogaster. Sci. Rep. 6, 20259. doi:10.1038/srep20259.

Khoobdel, M., Dehghan, H., Dayer, M. S., Asadi, A., Sobati, H., and Yusuf, M. A. (2021). Evaluation of a newly modified eight-chamber-olfactometer for attracting German cockroaches Blattella germanica (Dictyoptera: Blattellidae). Int. J. Trop. Insect Sci. 41, 979–989. doi:10.1007/s42690-020-00279-5.

Krofczik, S., Menzel, R., and Nawrot, M. P. (2008). Rapid odor processing in the honeybee antennal lobe network. Front. Comput. Neurosci. 2, 1–13. doi:10.3389/neuro.10.009.2008.

Lewis, L. P. C., Siju, K. P., Aso, Y., Friedrich, A. B., Bulteel, A. J. B., Rubin, G. M., et al. (2015). A higher brain circuit for immediate integration of conflicting sensory information in Drosophila. Curr. Biol. 25, 2203–2214. doi:10.1016/j.cub.2015.07.015.

Li, F., Lindsey, J. W., Marin, E. C., Otto, N., Dreher, M., Dempsey, G., et al. (2020). The connectome of the adult Drosophila mushroom body provides insights into function. Elife 9, 1–86. doi:10.7554/eLife.62576.

Li, Y., and Strausfeld, N. J. (1997). Morphology and sensory modality of mushroom body extrinsic neurons in the brain of the cockroach, Periplaneta americana. J. Comp. Neurol. 387, 631–650. doi:10.1002/(SICI)1096-9861(19971103)387:4<631::AID-CNE9>3.0.CO;2-3.

Li, Y., and Strausfeld, N. J. (1999). Multimodal efferent and recurrent neurons in the medial lobes of cockroach mushroom bodies. J. Comp. Neurol. 409, 647–663. doi:10.1002/(SICI)1096-9861(19990712)409:4<647::AID-CNE9>3.0.CO;2-3.

Meier, R., Egert, U., Aertsen, A., and Nawrot, M. P. (2008). FIND — A unified framework for neural data analysis. Neural Networks 21, 1085–1093. doi:10.1016/j.neunet.2008.06.019.

Menzel, R. (2012). The honeybee as a model for understanding the basis of cognition. Nat. Rev. Neurosci. 13, 758–768. doi:10.1038/nrn3357.

Mizunami, M., Okada, R., Li, Y., and Strausfeld, N. J. (1998). Mushroom bodies of the cockroach: activity and identities of neurons recorded in freely moving animals. J. Comp. Neurol. 402, 501–519. doi:10.1002/(SICI)1096-9861(19981228)402:4<501::AID-CNE5>3.0.CO;2-M.

Mote, M. I., and Goldsmith, T. H. (1970). Spectral sensitivities of color receptors in the compound eye of the cockroach Periplaneta. J. Exp. Zool. 173, 137–145. doi:10.1002/jez.1401730203.

Nawrot, M. P., Aertsen, A., and Rotter, S. (1999). Single-trial estimation of neuronal firing rates: From single-neuron spike trains to population activity. J. Neurosci. Methods 94, 81–92. doi:10.1016/S0165-0270(99)00127-2.

Okada, R., Ikeda, J., and Mizunami, M. (1999). Sensory responses and movement-related activities in extrinsic neurons of the cockroach mushroom bodies. J. Comp. Physiol. A 185, 115–129. doi:10.1007/s003590050371.

Okada, R., Sakura, M., and Mizunami, M. (2003). Distribution of dendrites of descending neurons and its implications for the basic organization of the cockroach brain. J. Comp. Neurol. 458, 158–174. doi:10.1002/cne.10580.

Owald, D., Felsenberg, J., Talbot, C. B., Das, G., Perisse, E., Huetteroth, W., et al. (2015). Activity of defined mushroom body output neurons underlies learned olfactory behavior in Drosophila. Neuron 86, 417–427. doi:10.1016/j.neuron.2015.03.025.

Owald, D., and Waddell, S. (2015). Olfactory learning skews mushroom body output pathways to steer behavioral choice in Drosophila. Curr. Opin. Neurobiol. 35, 178–184. doi:10.1016/j.conb.2015.10.002.

Paoli, M., Nishino, H., Couzin-Fuchs, E., and Galizia, C. G. (2020). Coding of odour and space in the hemimetabolous insect Periplaneta americana. J. Exp. Biol. 223, 1–14. doi:10.1242/jeb.218032.

Pedregosa, F., Varoquaux, G., Gramfort, A., Michel, V., Thirion, B., Grisel, O., et al. (2012). Scikit-learn: machine learning in Python. J. Mach. Learn. Res. 12, 2825–2830. Available at: http://arxiv.org/abs/1201.0490.

Rickert, J., Riehle, A., Aertsen, A., Rotter, S., and Nawrot, M. P. (2009). Dynamic encoding of movement direction in motor cortical neurons. J. Neurosci. 29, 13870–13882. doi:10.1523/JNEUROSCI.5441-08.2009.

Sakura, M., Okada, R., and Mizunami, M. (2002). Olfactory discrimination of structurally similar alcohols by cockroaches. J. Comp. Physiol. A 188, 787–797. doi:10.1007/s00359-002-0366-y.

Sayin, S., De Backer, J.-F., Siju, K. P., Wosniack, M. E., Lewis, L. P., Frisch, L.-M., et al. (2019). A neural circuit arbitrates between persistence and withdrawal in hungry Drosophila. Neuron 104, 544–558. doi:10.1016/j.neuron.2019.07.028.

Schubert, M., Hansson, B. S., and Sachse, S. (2014). The banana code—natural blend processing in the olfactory circuitry of Drosophila melanogaster. Front. Physiol. 5, 1–13. doi:10.3389/fphys.2014.00059.

Shiraiwa, T., and Carlson, J. R. (2007). Proboscis Extension Response (PER) Assay in Drosophila. J. Vis. Exp., 2–3. doi:10.3791/193.

Siju, K. P., Štih, V., Aimon, S., Gjorgjieva, J., Portugues, R., and Grunwald Kadow, I. C. (2020). Valence and state-dependent population coding in dopaminergic neurons in the fly mushroom body. Curr. Biol. 30, 2104–2115. doi:10.1016/j.cub.2020.04.037.

Strausfeld, N. J., Sinakevitch, I., Brown, S. M., and Farris, S. M. (2009). Ground plan of the insect mushroom body: functional and evolutionary implications. J. Comp. Neurol. 513, 265–291. doi:10.1002/cne.21948.

Strube-Bloss, M. F., Herrera-Valdez, M. A., and Smith, B. H. (2012). Ensemble response in mushroom body output neurons of the honey bee outpaces spatiotemporal odor processing two synapses earlier in the antennal lobe. PLoS One 7, 1–13. doi:10.1371/journal.pone.0050322.

Strube-Bloss, M. F., Nawrot, M. P., and Menzel, R. (2011). Mushroom body output neurons encode odor reward associations. J. Neurosci. 31, 3129–3140. doi:10.1523/JNEUROSCI.2583-10.2011.

Strube-Bloss, M. F., Nawrot, M. P., and Menzel, R. (2016). Neural correlates of side-specific odour memory in mushroom body output neurons. Proc. R. Soc. B 283, 20161270. doi:10.1098/rspb.2016.1270.

Strube-Bloss, M. F., and Rössler, W. (2018). Multimodal integration and stimulus categorization in putative mushroom body output neurons of the honeybee. R. Soc. Open Sci. 5, 171785. doi:10.1098/rsos.171785.

Tsao, C.-H., Chen, C.-C., Lin, C.-H., Yang, H.-Y., and Lin, S. (2018). Drosophila mushroom bodies integrate hunger and satiety signals to control innate food-seeking behavior. Elife 7, 1–35. doi:10.7554/eLife.35264.

Tully, T., and Quinn, W. G. (1985). Classical conditioning and retention in normal and mutant Drosophila melanogaster. J. Comp. Physiol. A 157, 263–277. doi:10.1007/BF01350033.

Walther, J. B. (1958). Changes induced in spectral sensitivity and form of retinal action potential of the cockroach eye by selective adaptation. J. Insect Physiol. 2, 142–151. doi:10.1016/0022-1910(58)90038-6.

Yagi, R., Mabuchi, Y., Mizunami, M., and Tanaka, N. K. (2016). Convergence of multimodal sensory pathways to the mushroom body calyx in Drosophila melanogaster. Sci. Rep. 6, 1–8. doi:10.1038/srep29481.

Yetman, S., and Pollack, G. S. (1987). Proboscis extension in the blowfly: directional responses to stimulation of identified chemosensitive hairs. J. Comp. Physiol. A 160, 367–374. doi:10.1007/BF00613026.

